# Fast and functionally specific cortical thickness changes induced by visual stimulation

**DOI:** 10.1101/2022.02.25.482013

**Authors:** Natalia Zaretskaya, Erik Fink, Ana Arsenovic, Anja Ischebeck

**Affiliations:** Department of Cognitive Psychology and Neuroscience, Institute of Psychology, University of Graz, Austria; BioTechMed-Graz, Austria

**Keywords:** cortex, MRI, plasticity, thickness, V1

## Abstract

Structural characteristics of the human brain serve as important markers of brain development, aging, disease progression and neural plasticity. They are considered stable properties, changing slowly over time. Multiple recent studies reported that structural brain changes measured with MRI may occur much faster than previously thought, within hours or even minutes. The mechanisms behind such fast changes remain unclear, with hemodynamics as one possible explanation. Here we investigated the functional specificity of cortical thickness changes induced by a flickering checkerboard and compared the them to BOLD fMRI activity. We found that checkerboard stimulation led to a significant thickness increase, which was driven by an expansion at the gray-white matter boundary, functionally specific to V1, confined to the retinotopic representation of the checkerboard stimulus, and amounted to 1.3 % or 0.022 mm. Although functional specificity and the effect size of these changes were comparable to those of the BOLD signal in V1, thickness effects were substantially weaker in V3. Furthermore, a comparison of predicted and measured thickness changes for different stimulus timings suggested a slow increase of thickness over time, speaking against a hemodynamic explanation. Altogether, our findings suggest that visual stimulation can induce structural gray matter enlargement measurable with MRI.

## Introduction

Quantification of the structural characteristics of the human brain using MRI scans is central for various fields of neuroscience. For example, gray matter changes serve as an important biomarker of neurodegenerative diseases and their progression (Suri et al. 2014; Sterling et al. 2016; Mazón et al. 2018). It is also widely used to study brain changes that accompany psychiatric diseases (Fusar-Poli and Meyer-Lindenberg 2016) and neurodevelopmental disorders (Wee et al. 2014) in search for suitable biomarkers. Anatomical brain parameters are central in studies of healthy brain development (Oishi et al. 2013) and aging (MacDonald and Pike 2021). Finally, structural brain changes are critical for the studies of experience-dependent plasticity (Lövdén et al. 2013; Draganski et al. 2014; Calmels 2020), as well as reorganization after stroke (Zatorre et al. 2012; Sampaio-Baptista et al. 2018).

In such quantitative morphometry studies, structural brain characteristics are typically viewed as rather stable trait-like properties that change slowly over time, with typical time scales ranging from days to years or even decades. However, recent human MRI studies demonstrate that measurable structural brain changes may occur much faster than previously thought. For example, reduction of the gray matter volume can be observed throughout one day, reflecting possible influences of the circadian rhythm on structural brain characteristics (Trefler et al. 2016). Structural gray matter changes can even be detected within a few hours after different forms of learning (Taubert et al. 2016; Hofstetter et al. 2017; Brodt et al. 2018). A recently published study demonstrated gray matter reduction after as little as two minutes of task (Olivo et al. 2022). The fast nature of these effects raises the question about their underlying mechanisms. Specifically, it is not clear whether these fast changes reflect the true structural changes of neural tissue observed in animal studies (Holtmaat and Svoboda 2009).

The exact cellular mechanisms that drive morphological gray matter changes observed in human MRI may be quite diverse. Cortical volume reduction in neurodegenerative diseases and in healthy aging is thought to be, at least to some extent, a consequence of true loss of gray matter tissue (Esiri 2007; Wyss-Coray 2016). Mechanisms behind plasticity-dependent gray matter *increases* are far less understood. Outside of brain regions known to show neurogenesis in adulthood (olfactory bulb, dentate gyrus of the hippocampus), gray matter changes are typically attributed to mechanisms such as axonal sprouting, dendritic branching, dendritic spine formation or gliogenesis (Wenger et al. 2017). Crucially, several prior studies indicated that the MRI-derived apparent tissue changes may be an artifact produced by hemodynamics. The T1 value of the arterial blood is close to that of gray matter at 3 T magnetic field strength (Lu et al. 2004). Therefore, blood flow change within the gray matter may manifest itself as image intensity change and, consequently, as gray matter volume change (Franklin et al. 2013; Ge et al. 2017). Furthermore, an increase of the cerebral blood volume due to vessel dilation, which also accompanies ongoing neural activity, may contribute to the gray matter tissue expansion and consequently, to cortical thickness increase. While true structural changes may happen at various time scales, but are typically considered slow, hemodynamic changes are nearly instantaneous relative to the timescale of a typical structural scan duration of several minutes. Thus, hemodynamics may account for the fast structural changes observed in MRI.

The majority of previous studies examining fast gray matter changes focused on learning-dependent plasticity paradigms, in which structural changes were attributed to the learned skill (Taubert et al. 2016; Hofstetter et al. 2017; Brodt et al. 2018). These studies demonstrated some degree of functional specificity, with primary effects confined to brain regions involved in skill acquisition. The evidence for gray matter changes driven by factors other than learning remains scarce. In one previous study, changes induced by visual stimulation could be detected on a time scale as short as several minutes (Månsson et al. 2020). Surprisingly, the main effects in this study were observed outside of the primary visual cortex, within the lingual and fusiform gyri. Therefore, it remains unclear to what degree visual stimulation effects are functionally specific and how their functional specificity compares to hemodynamics. In addition, since in this study emotionally arousing images were compared with a blank screen, the observed changes could have, in principle, been driven by both the bottom-up sensory stimulation as well as top-down processing, such as attention.

In the current study, we investigated the bottom-up sensory and top-down attentional contribution to short-term cortical thickness changes. Furthermore, we investigated the functional specificity of short-term thickness changes induced by visual stimulation by examining the exact spatial pattern of changes across the visual cortex and comparing the structural effects with the classical T2*-weighted functional MRI.

## Methods

### Data and code availability statement

All analysis scripts, ROI data and group fsaverage MRI results are available at osf.io ([link will be provided upon manuscript acceptance]). Our institution’s data protection policy currently prohibits us from publicly sharing native space MRI data. However, it can be shared with other researchers upon direct requests addressed to the corresponding author.

### Participants

Sixty healthy adult volunteers participated in the experiment. Data of three participants had to be excluded due to incomplete dataset (2 participants) or failure of automatic cortical surface reconstruction (1 participant). This left 57 participants (mean age: 24.2, *SD*: 3.2, 35 female). All participants had normal or corrected-to-normal vision (≤ -0.5 diopters) and no history of neurological impairments. Subjects were screened for MRI-contraindications and had signed a written informed consent. The study was conducted according to the Declaration of Helsinki and was approved by the ethics committee of the University of Graz.

### Stimulus and experimental procedures

Visual stimulation was generated using MATLAB 2019b (MathWorks, Natick, MA) with the Psychophysics Toolbox 3 extensions (Brainard 1997; Pelli 1997) on a Linux Ubuntu 18.04 LTS computer and presented using a linearized MR-compatible screen (Nordic NeuroLab, Bergen, Norway) placed behind the scanner bore (1920×1080 resolution, 60 Hz). The screen was viewed through a mirror mounted onto the RF coil at a total distance of 143.5 cm, subtended 27.72°×15.79° in visual angle units and had a maximum luminance of 405 cd/m^2^.

The stimulus layout consisted of a central fixation dot (diameter: 6.02° of visual angle), the surrounding central gray disk (diameter: 8.62°), the outer contrast-reversing checkerboard pattern and two letter streams presented simultaneously on both sides of the screen (Figure 1). The letter circles had a diameter of 4.81° and were placed at an eccentricity of 10° from the fixation point. New letters appeared continuously throughout each run at a frequency of 2 Hz (i.e., new letter every 500 ms). The onset of new letters was synchronized between hemifields, but the letter identity and color were randomized independently for each hemifield.

**Figure 1.**
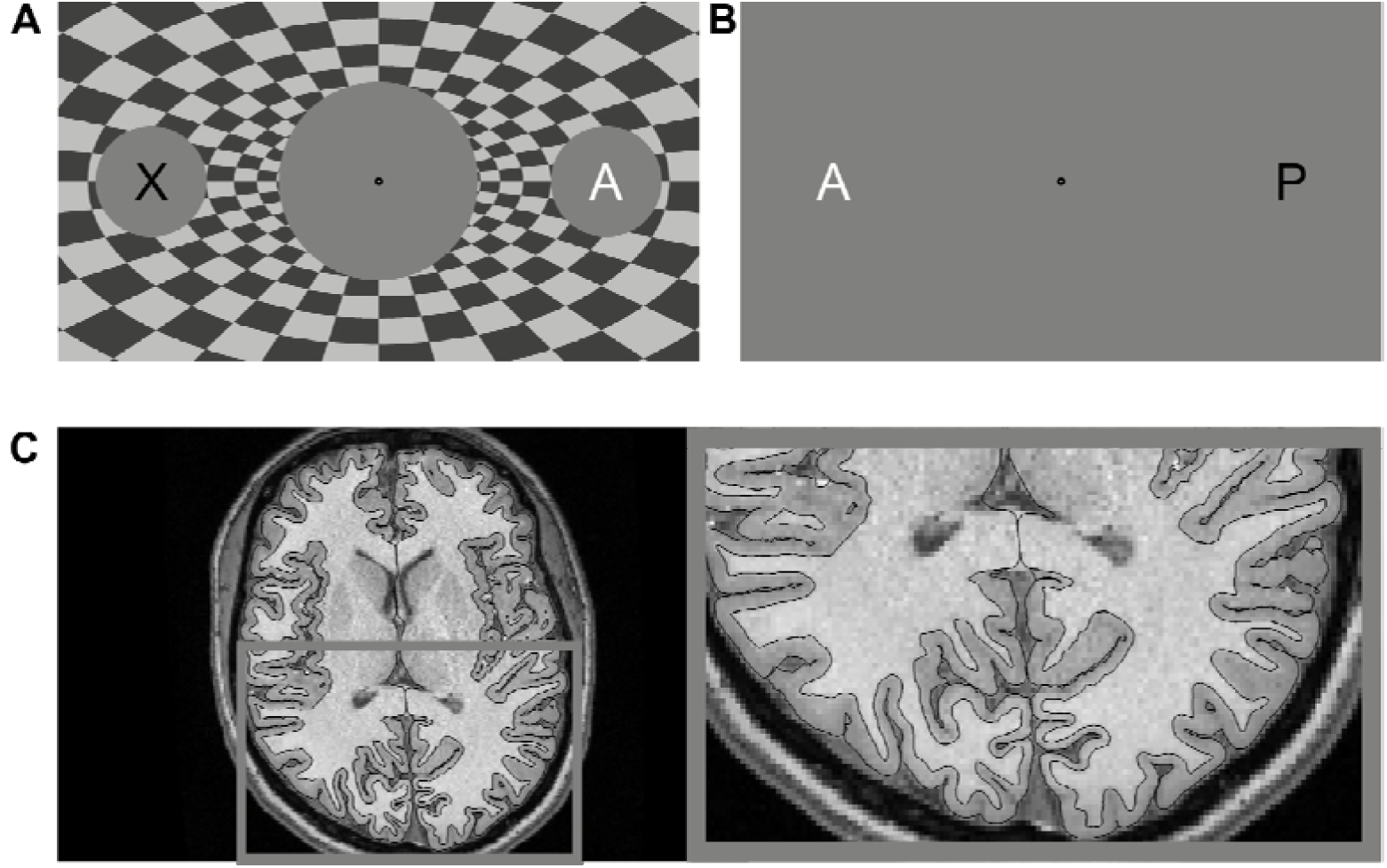
Visual stimulus and data analysis. The stimulus consisted of either flickering checkerboard (A) or gray screen (B) with a central fixation dot surrounded by a gray disk. The two letter streams surrounded by gray disks were presented on both sides simultaneously. The letters could either be black or white. Depending on the experimental run, participant’s task was either to detect white letters irrespective of identity (color task), or a white letter « A » (letter task). (C) Axial slice of one MPRAGE volume acquired during one run (checkerboard stimulation, color task) of a representative participant along with reconstructed cortical surfaces used for thickness estimation. Gray square indicates area around the occipital lobe that is shown on the right to illustrate the surface reconstruction quality.

The main experiment consisted of four structural MRI runs. We used a 2×2 factorial experimental design with factors “stimulation” (on, off), and “task” (color task, letter task), yielding four experimental conditions. In the color task, participants were required to press a button whenever they saw a white letter on either side of the fixation point. In the letter task, participants were required to press a button whenever they saw the letter “A” in white color on either side of the fixation point. The target probability of 2.5% was identical in both tasks. Each condition started synchronously with the onset of the structural scan acquisition and was presented for the whole duration of 7 minutes, with the order of conditions pseudorandomized and counterbalanced across participants.

To compare the spatial pattern of thickness changes with the spatial pattern of BOLD signal changes observed in fMRI, we conducted an additional fMRI experiment within the same session. The fMRI experiment had an identical stimulus layout and task, but different timings. The functional runs consisted of 12-second trials with either checkerboard present or absent while the letter streams were always on, interleaved with fixation baseline consisting of the gray background only (without the letter streams). Each task was presented in a separate functional run, with the order of tasks counterbalanced across participants. A total of two runs, one for the color and one for the letter task, were acquired. The functional scans were preceded by a separate standard T1-weighted structural scan, which was used exclusively for the surface-based fMRI analysis. The functional experiment preceded the main experiment and was separated from the latter by a 15-minute-long diffusion acquisition (without any task), which was not related to the goals of this experiment.

### MRI data acquisition

Experiments were performed on a 3 T whole-body MRI scanner (Magnetom Vida, Siemens Healthineers, Erlangen, Germany) with a 64-channel Head/Neck coil (Siemens Healthineers, Erlangen, Germany). The acquisition parameters for each of the protocols are listed in Table 1.

**Table 1.** Protocol overview

| **Protocol type** | **MPRAGE (fMRI analysis)** | **2D GE-EPI** | **MPRAGE (main experiment)** |
| --- | --- | --- | --- |
| **Isotropic voxel size (mm)** | 1 | 3 | 0.6 |
| **TR (ms)** | 2530 | 3220 | 2510 |
| **TE (ms)** | 3.88 | 32 | 4.1 |
| **TI (ms)** | 1200 | n/a | 1200 |
| **FA** | 7° | 82° | 7° |
| **matrix** | 256×256 | 76×76 | 416×416 |
| **slices** | 176 | 46 | 224 |
| **PE** | A>>P | A>>P | R>>L |
| **acceleration** | GRAPPA 2 | none | GRAPPA 2 |
| **duration (min.)** | 6:03 | 8:22 | 7:11 |
| **partial Fourier** | Off | Off | Off |
| **bandwidth (Hz/pixel)** | 190 | 1828 | 200 |

### Data analysis

#### Structural MRI analysis

High-resolution structural scans from the main experiment with visual stimulation (see table 1, last column) underwent the following preprocessing steps. First, the reduced image intensity of the top and bottom slices, which is due to the slab-selective acquisition, was corrected using the SPM12 implementation of the unified segmentation algorithm (Ashburner and Friston 2005). After this, cortical thickness estimation for each volume was performed with FreeSurfer 7.1.0 (Fischl and Dale 2000a) within the longitudinal processing pipeline (Reuter and Fischl 2011; Reuter et al. 2012), which included several steps. As the first step, each of the four structural volumes was processed using the FreeSurfer’s standard recon-all stream. The major processing steps of the standard recon-all include brain mask creation, bias correction, segmentation of gray and white matter and creation of the two boundary surfaces, one at the intersection of the gray and white matter (termed “white surface”) and one at the intersection of the gray matter and cerebrospinal fluid (termed “pial surface”). Cortical thickness is determined from the two surfaces and the distance between the white and the pial surface at each vertex location along the surface normal. As the second step of the longitudinal processing pipeline, all four individual volumes were used to create a subject-specific unbiased template volume, termed “base”, which represents the average anatomy of each subject (Reuter et al. 2010). The base was also processed with FreeSufer’s recon-all stream and the information from the base (including the location of the cortical surfaces) was then used as an initial guess for the surfaces of individual time points, which are adjusted based on the image information of the individual volumes. The advantages of longitudinal stream include the unbiased and identical processing of every time point as well as the correspondence of surface vertices (Reuter and Fischl 2011; Reuter et al. 2012). Vertex correspondence between individual time points and the base additionally allowed us to determine identical surface locations in every volume and measure the displacement of individual surfaces (for details see below).

Although our native resolution was 0.6 mm isotropic, we used FreeSufer’s default setting of downsampling each volume to 1 mm isotropic voxel size. We decided against processing the data at native resolution for the following reasons. First, due to partial volume modelling, FreeSufer is sensitive to thickness changes that are much smaller than a typical 1 mm voxel size (Fischl and Dale 2000b). Previous assessments of test-retest reliability indicated a variation of FreeSurfer-derived thickness estimates of 0.05 mm (1.9 %) or less (Wang et al. 2008) or even 0.03 mm or less (Han et al. 2006), which enables it to detect very small changes. Second, our individual 0.6 mm isotropic resolution at 3 T had relatively low SNR compared to typical 1 mm^3^ scans. Interpolation involved in the downsampling step therefore helped to increase the image SNR. Third, we have previously shown that higher resolution increases the accuracy (i.e., “correctness”) of the surface placement (Zaretskaya et al. 2018), but there is no evidence that it also helps to increase the precision (i.e., the scan-to-scan consistency), which is more important for detecting change. Finally, processing the data at 1 mm^3^ was computationally more efficient.

##### Whole-brain thickness analysis

Cortical thickness maps derived from each subject and each run were then projected onto the FreeSurfer fsaverage standard surface space and smoothed with a 10 mm full width at half maximum (FWHM) 2D Gaussian kernel. Smoothing in neuroimaging is criticized for reducing the experimental effect size and localization of cortical areas, especially in the case of volumetric analysis (Stelzer et al. 2014; Coalson et al. 2018). In the worst case, it can entirely cancel an effect that exists on a very fine spatial scale, like the ocular dominance column signal (Zaretskaya et al. 2020). This is why smoothing in the current study was performed exclusively for the whole-brain analysis. In the whole-bran analysis smoothing is necessary to meet the assumptions of multiple comparison correction procedures and may also be useful for increasing the SNR of noisy data. We have chosen the kernel width of 10 mm following the study of Månsson et al. (2020). Differences between conditions were assessed at each surface vertex by modelling the main effects of checkerboard and task as well as their interaction. Correction for multiple comparisons was performed using a parametric Gaussian-based cluster-level inference with a cluster-forming threshold corresponding to p < 0.05 (two-tailed test), a cluster-wise threshold of p < 0.05 (two-tailed test), and an additional Bonferroni correction of the cluster-wise significance for two hemispheres.

##### Region of interest analysis

To further determine the functional specificity of the cortical thickness changes, we conducted a region of interest (ROI) analysis within the early visual cortex. As there was no effect of task in the whole-brain analysis (see “Results” section for details), we focused primarily on the stimulation effect, averaging values of the two tasks. First, we predicted surface location and retinotopic organization (eccentricity and polar angle) of the early visual areas (V1, V2 and V3) in each subject using the retinotopic surface template by Benson et al. (Benson et al. 2012, 2014)^1^. The eccentricity information of the template was subsequently used to divide the early visual areas into three eccentricities: center (corresponding to the gray disk), checkerboard representation, and periphery (eccentricities outside of the screen). This resulted in three ROIs per subject and hemisphere (also illustrated in Figure 3A). Native space thickness values for the “stimulation on” and “stimulation off” conditions (averaged over tasks) were averaged over all vertices within each ROI. Our ROI analysis is based exclusively on unsmoothed vertex values.

**Figure 3.**
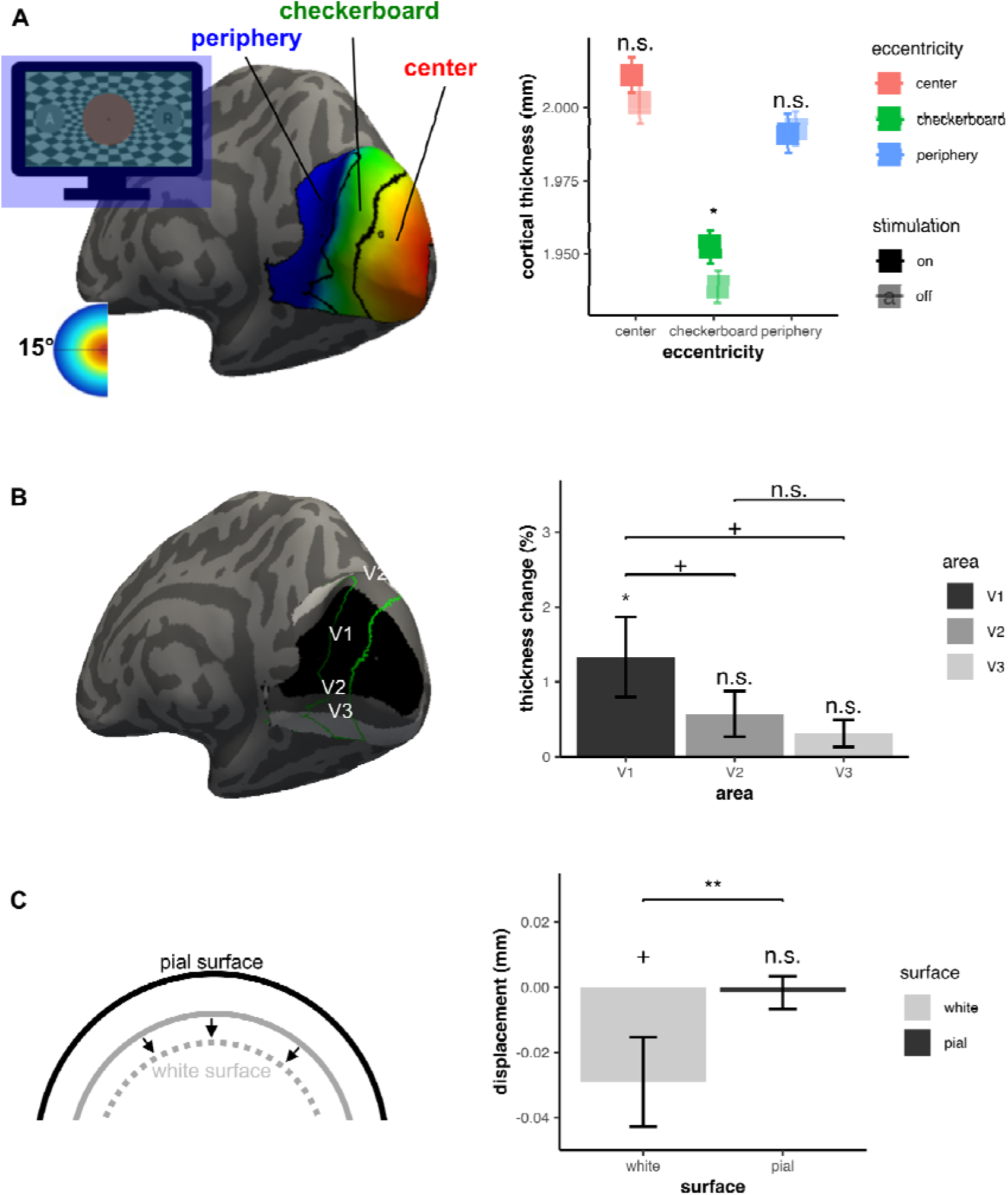
Specificity of cortical thickness change across the early visual cortex. A) Spatial specificity. Stimulus layout and the representation of the three eccentricities on the inflated cortical surface of a representative participant (left), as well as the average cortical thickness measured within each eccentricity for the “stimulation on” and “stimulation off” conditions (right). Only the checkerboard eccentricity demonstrates significant effect of stimulus presence. B) Functional specificity. Location of the visual areas V1, V2, V3 for the same participant and the outline of the checkerboard eccentricity (left), as well as the cortical thickness change induced by the stimulation (right). Only the checkerboard representation of V1 showed a significant change of thickness due to stimulation. C) Schematic representation of the surface displacement (left) and the actual measured displacement (right) of the white and pial boundary surfaces. The white surface moved significantly further inwards compared to the pial surface, which produced a net cortical thickness increase within the checkerboard representation of V1. *p < 0.05 corrected, +p < 0.05 uncorrected; error bars indicate SEM.

In addition, to determine whether thickness changes were driven by the displacement of the gray-white matter boundary (demarcated by the position of the white surface), the gray-CSF boundary (demarcated by the position of the pial surface), or by both, we analyzed the change in position of each surface separately. This was done following the procedure described in (Zaretskaya et al. 2018). Due to our usage of the longitudinal pipeline, there is a natural vertex correspondence between the surfaces of each of the four brain volumes and the surfaces of the subject’s template (“base”). Therefore, we could calculate the average displacement between the surface position during the “stimulation on” and the average position during the “stimulation off” conditions along the surface normal at every vertex location. By FreeSurfer convention, normals are pointing outwards, i.e., from the white matter to the gray matter. Therefore, positive displacement corresponds to the outward surface movement and negative displacement corresponds to the inward surface movement. Surface normals of the base template were used as a reference to calculate the displacement sign. Displacement values were averaged within each ROI for the subsequent statistical analysis.

##### Statistical hypotheses tests

All statistical analyses were performed with RStudio 2021.09.1 (RStudio Team 2021). In cases where an ANOVA was performed, Greenhouse-Geisser correction for non-sphericity was automatically applied to within-subject factors where Mauchly’s test of sphericity was significant at p < 0.05.

To determine whether visual stimulation effects on cortical thickness were specific to the eccentricity representing the checkerboard stimulus, ROI values were subjected to a three-way repeated measures ANOVA with factors “hemisphere” (left, right), “eccentricity” (center, checkerboard, periphery) and “stimulation” (on, off). The hemisphere factor was included to additionally test for a potential lateralization of the effect, which appeared to be more pronounced in the right hemisphere in the whole-brain analysis.

To determine whether the observed effects within the central eccentricity representing the checkerboard are present throughout all three early visual areas or are confined to the primary visual cortex (V1), we further split the checkerboard eccentricity ROI into three subparts overlapping V1, V2 and V3 (as demonstrated in Figure 3B). We quantified the stimulation effect in each area as the difference of thickness during stimulation on and off screen, divided by the average thickness of both and multiplied by 100. We then performed a one-way repeated measures ANOVA of the stimulation effect across the three areas. Since no effect of hemisphere and no interaction between hemisphere and checkerboard factors was found in the first analysis, values from both hemispheres were averaged. In addition, since one-way ANOVA only compared areas with each other, we also tested for the presence of an effect in each area by comparing the stimulation effect with zero using a two-tailed one-sample t-test. For determining the displacement of each surface, the displacement of white and pial surface was compared against zero as well as with each other using a one-sample and a paired-sample two-tailed t-test, respectively.

##### Control analysis of image intensity

To rule out that the cortical thickness changes are due to changes in intracortical image intensity, which could be caused by hemodynamic changes, we repeated our three-way repeated measures ANOVA ROI analysis, but with “percent gray-white contrast” as the dependent measure instead of thickness. To calculate the gray-white contrast, white matter intensity is sampled 1 mm below the white surface and the gray matter intensity is sampled at 30% of the cortical thickness above the white surface along the surface normal at every vertex location. Percent gray-white contrast is defined as the difference between the white and gray matter intensities, divided by their average and multiplied by 100. If these values were calculated based on a single pair of surfaces derived from the subject’s template, changes in cortical thickness of individual volumes would have led to a misalignment of the template and the individual gray-white boundary, leading to sampling at the “wrong” depths along the surface normal (for example, if the cortex of an individual volume is thicker than that of a template, the more superficial white matter locations in the template may correspond to deeper gray matter in that volume). Therefore, for this analysis percent gray-white contrast values were derived from each volume’s own pair of surfaces that are generated within the longitudinal stream. Because of this, displacement of one or both surfaces that are associated with changes in cortical thickness should not bias or compromise our analysis.

#### Functional MRI analysis

Functional data were preprocessed using fMRIprep version 20.2.3 (Esteban et al. 2019) and included motion correction, slice-time correction, distortion correction using an additional scan acquired with opposite phase-encoding direction, coregistration with the 1 mm MPRAGE structural scan acquired in the beginning of the experiment (see Table 1, 1^st^ column), and projection onto the individual subject’s reconstructed cortical surface as well as onto the FreeSurfer fsaverage template. Using an independent structural scan for surface-based fMRI analysis allowed us to ensure full independence of our fMRI and thickness effects. For the whole-brain group analysis, fsaverage surface-space data were smoothed with a 2D Gaussian kernel of 5 mm FWHM. No smoothing was applied in the individual-space ROI analysis. Functional data analysis was subsequently performed using the FreeSurfer functional analysis stream (FS-FAST) and a general linear model (GLM) approach. Four conditions of interest (stimulation on and off during the color task, stimulation on and off during the letter task) were convolved with a standard hemodynamic response function and entered into a GLM together with nuisance regressors modelling slow scanner drifts. Stimulation effects were assessed using the contrasts “stimulation on vs. off”. For the whole-brain group analysis, individual contrasts were entered into a second level GLM testing for non-zero effects using weighted least squares approach (FreeSurfer’s *mri_glmfit* with “-wls” flag). For the ROI analysis, individual contrast values were additionally converted into percent contrast. The latter was defined as the difference between the stimulation on and stimulation off regressors, divided by the model offset and multiplied by 100.

#### Comparison of structural and functional effects

To further determine how the observed thickness changes relate to the measured BOLD fMRI effects, we conducted several additional analyses. First, following a procedure similar to our main thickness analysis, we identified vertices representing different eccentricities and areas V1, V2, and V3 on the structural scans used for functional analysis, and averaged the BOLD stimulation effect (defined as the “stimulation on vs. off”, divided by the GLM model offset and multiplied by 100) within the checkerboard representations of areas V1, V2 and V3. Furthermore, to compare the strength of the two effects (thickness and BOLD), we performed a paired t-test for each area, correcting for 3 comparisons with Bonferroni method. Finally, to compare the functional specificity of the two effects, we further subdivided all V1 eccentricities between 0 and 30° into bins of 1°, averaged effects across vertices within each bin, and compared the two effects using a non-parametric permutation test with cluster-wise inference (Maris and Oostenveld 2007) using the MATLAB permutest.m function by E. M. Gerber^2^.

Because spatial overlap between the two effects does not imply a relationship between them, we conducted two additional analyses directly aimed at testing for a relationship between the thickness change and the BOLD effect. First, we calculated Pearson’s correlation coefficient between the thickness effect and the BOLD effect within the checkerboard representation of V1 across subjects, as well as within the two clusters that showed cortical thickness changes in the whole-brain analysis. A positive correlation would indicate an underlying relationship between the two effects. Second, we repeated a one-way ANOVA across areas with the thickness effect as a dependent variable, but additionally included the BOLD effect as a covariate. If the observed thickness change is indeed related to hemodynamics, including the BOLD signal as a covariate should abolish the thickness effect.

#### Exploratory analysis of the signal time course

To determine whether the observed effects are more similar to the relatively fast hemodynamic signal, or to a slow build-up more characteristic of a structural change, we conducted an additional exploratory analysis of the impact of condition order on the measured effect. If the signal we measure is related to hemodynamics, that is, it has a relatively steep rise within the structural scan acquisition period and remains constant throughout the stimulation/volume acquisition, it should be equally strong no matter in which order the checkerboard conditions are presented. If, however, the signal builds up slowly, that is, over the course of a volume acquisition, and takes the same amount of time to decay, certain condition sequences may be less optimal than others for detecting the stimulation effect in the cortical thickness data.

Simulations of the slow and fast effect build-up were performed in MATLAB 2019b. For every condition order we generated a time vector with 1 s resolution, and a stimulus function with a value of 1 whenever checkerboard was present on the screen and zero otherwise (Figure 5A). The short breaks (mean over subjects: 1.29 minutes, SD: 0.59) between the scans were assumed to have a zero duration. The predicted hemodynamic response was generated by convolving the stimulus function with a hemodynamic response function (first gamma function from the canonical SPM-HRF without the undershoot). The linear build-up was generated by gradually increasing the values during the checkerboard periods and decreasing the values during the baseline, until the signal reached 1 or 0, respectively. After this the signal was set to plateau. The *simulated* stimulation effect for each signal type and each condition order was quantified by calculating the difference between checkerboard and baseline values, divided by the mean of both values, and multiplying by 100. The *measured* stimulation effect (thickness change) for each condition order was plotted from the V1 representation of the checkerboard for comparison. We would like to emphasize that the measured thickness analysis per condition order is exploratory and should be regarded with caution, because there are only 9-10 observations per condition order (compared to 57 observations for the whole experiment).

**Figure 5.**
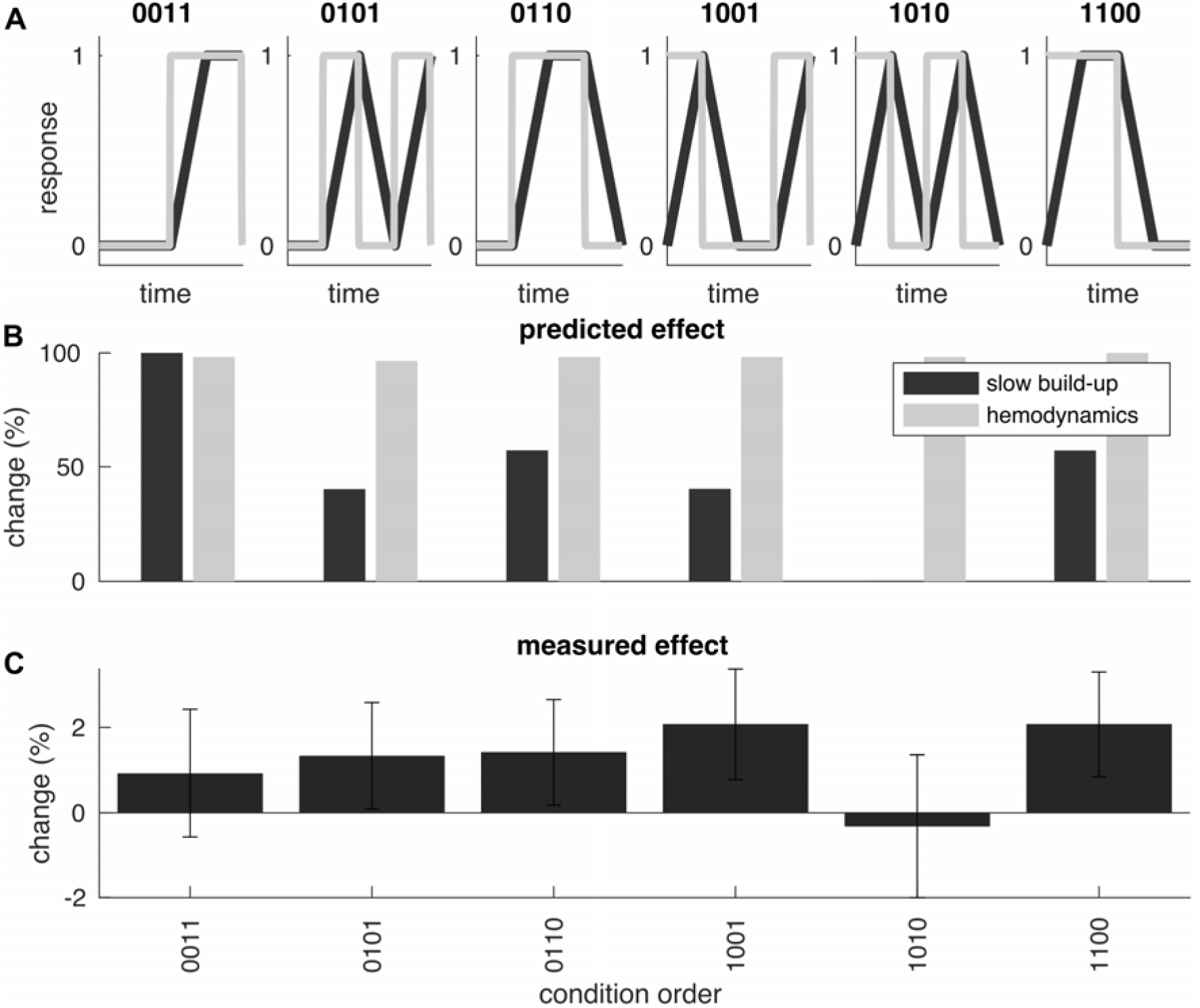
Impact of condition order on the measurable effect. A) Simulation of the hemodynamics (light gray) and slow linear increase (dark gray) of a response over time depending on the checkerboard condition order (0 corresponding to the gray baseline, while 1 corresponding to the checkerboard). B) Percent contrast change for each signal type calculated as difference divided by mean and multiplied by 100. Hemodynamic signal produces nearly equal responses for every condition order. Linear increase produces positive response in all but one condition combination (1010). C) Measured cortical thickness change in the checkerboard representation of V1, grouped by condition order. Similar to the slow buildup signal in the simulation, the effect is a nearly absent for the condition order 1010. Error bars represent SEM over subjects.

#### Behavioral data analysis

Behavioral performance of the participants was analyzed using a custom MATLAB script. Each button press recorded by the participants was classified as either a hit or a false alarm, depending on the presence of a target within an interval of 0 to 750 ms prior to the button press. For the main experiment, percentage of hits was calculated separately for each of the four conditions and submitted to a two-way repeated measures ANOVA with factors “stimulation” (on, off) and “task” (color, letter) using SPSS Statistics 28 (IBM Corporation, Armonk, NY, USA).

## Results

### Behavioral responses

The average hit rate in the task of the main experiment was 0.70 (SD=0.25). A two-way repeated measures ANOVA for the hit rates for the main experiment with factors “stimulation” (on, off) and “task” (color, letter) revealed no main effect of checkerboard (*F*(1, 56) = 2.086, *p* = 0.154, *η_p_*^2^ = 0.036), no main effect of task (*F*(1, 56) = 0.835, *p* = 0.365, *η_p_*^2^ = 0.015), and no interaction between checkerboard and task (*F*(1, 56) = 1.51, *p* = 0.225, *η_p_*^2^ = 0.026).

### Whole-brain thickness changes

We first assessed stimulation- and task-induced cortical thickness changes across the whole cortical surface. The vertex-wise GLM analysis revealed a significant main effect of checkerboard around the right calcarine sulcus, corresponding to higher cortical thickness values during checkerboard stimulation compared to gray screen (cluster size = 1268.01 mm^2^, Z = 3.50, cluster-level *p* = 0.0014, MNI coordinates: x = 12.3, y = -84.9, z = 0.1). In addition, there was a checkerboard-induced reduction in cortical thickness in the left superior frontal gyrus (cluster size = 1040.20 mm^2^, Z = -2.96, cluster-level *p* = 0.011, MNI coordinates: x = -15.0, y = 57.2, z = 16.4). The whole-brain thickness difference maps as well as the two significant clusters are illustrated in Figure 2 (see also supplementary Figure S1 for unsmoothed maps). Neither the main effect of task, nor the interaction between the task and stimulation revealed a significant effect anywhere across the cortical surface.

**Figure 2.**
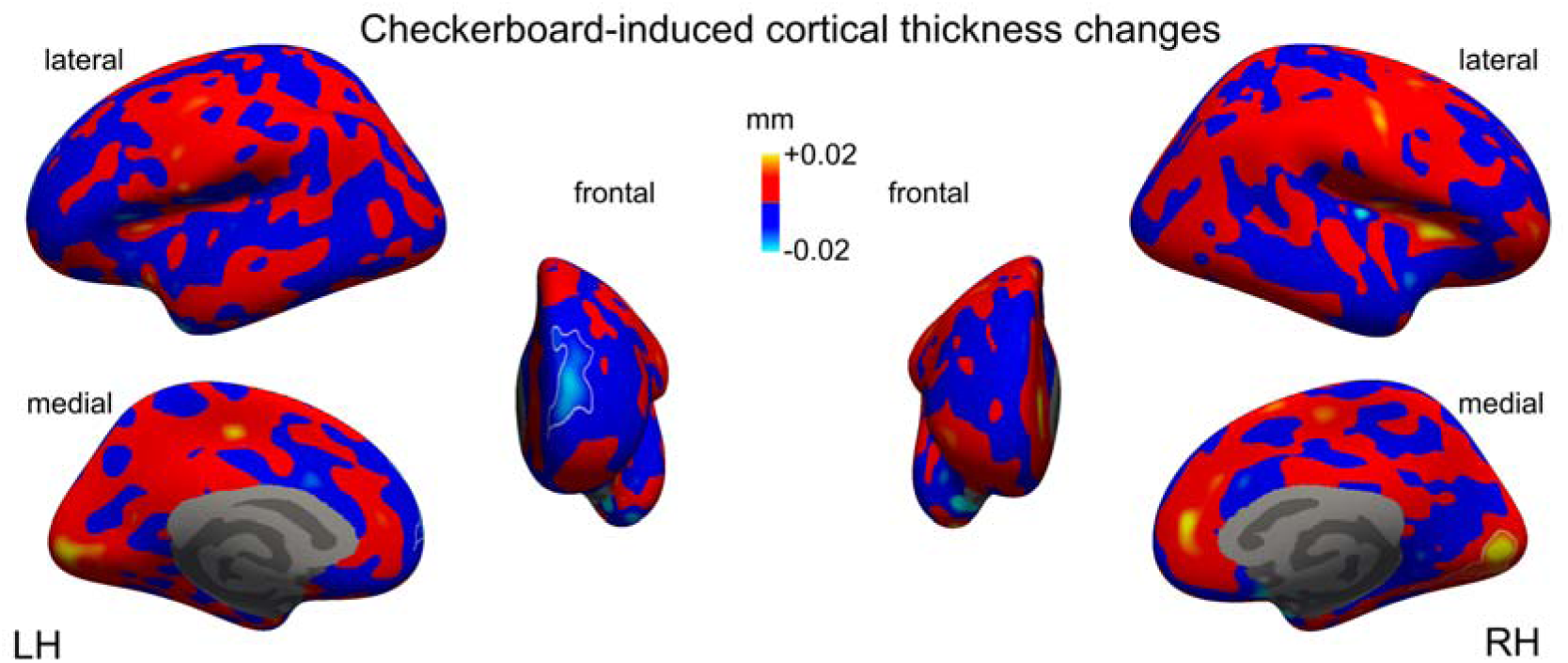
Whole-brain cortical thickness changes induced by checkerboard stimulation. Maps represent average difference between “stimulation on” and “stimulation off” conditions in mm at every surface vertex. The checkerboard effect appeared as thickness increase in the right calcarine sulcus (red-yellow colors) and as thickness decrease in the left superior frontal gyrus (blue-cyan colors). The two significant clusters (corrected for multiple comparisons at p<0.05, see methods) are shown as outlines. See also supplementary Figure S1 for unsmoothed maps.

### Topographic effects in the early visual cortex

Next, we tested whether cortical thickness increase occurred throughout the early visual areas or specifically within the topographic representation of the checkerboard stimulus and whether there were any hemisphere lateralization effects (e.g., significantly stronger effects in the right hemisphere, as suggested by the whole-brain group results). Figure 3 summarizes the results. Our three-way ANOVA revealed a significant effect of eccentricity (*F*(2,112) = 23.06, *p* < 0.001, *η_p_*^2^ = 0.29), a significant interaction between eccentricity and hemisphere (*F*(2,112) = 7.6, *p* = 0.008, *η_p_*^2^ = 0.12), and, crucially, a significant interaction between eccentricity and stimulation (*F*(1.55, 87.05) = 4.68, *p* = 0.02, *η_p_*^2^ = 0.08), suggesting topographic specificity of the checkerboard effect in area V1. All other main effects and interactions did not reach significance (full statistical results are provided in Table 2). Post-hoc paired-samples t-tests (averaged over hemispheres) revealed a significant checkerboard effect within the representation of the checkerboard (*t*(56) = 2.53, *p_bonf_* = 0.042), but not within the unstimulated center (*t*(56) = 1.69, *p_unc_*= 0.096), and not within the periphery (*t*(56) = -0.30, *p_unc_* = 0.770).

**Table 2.**
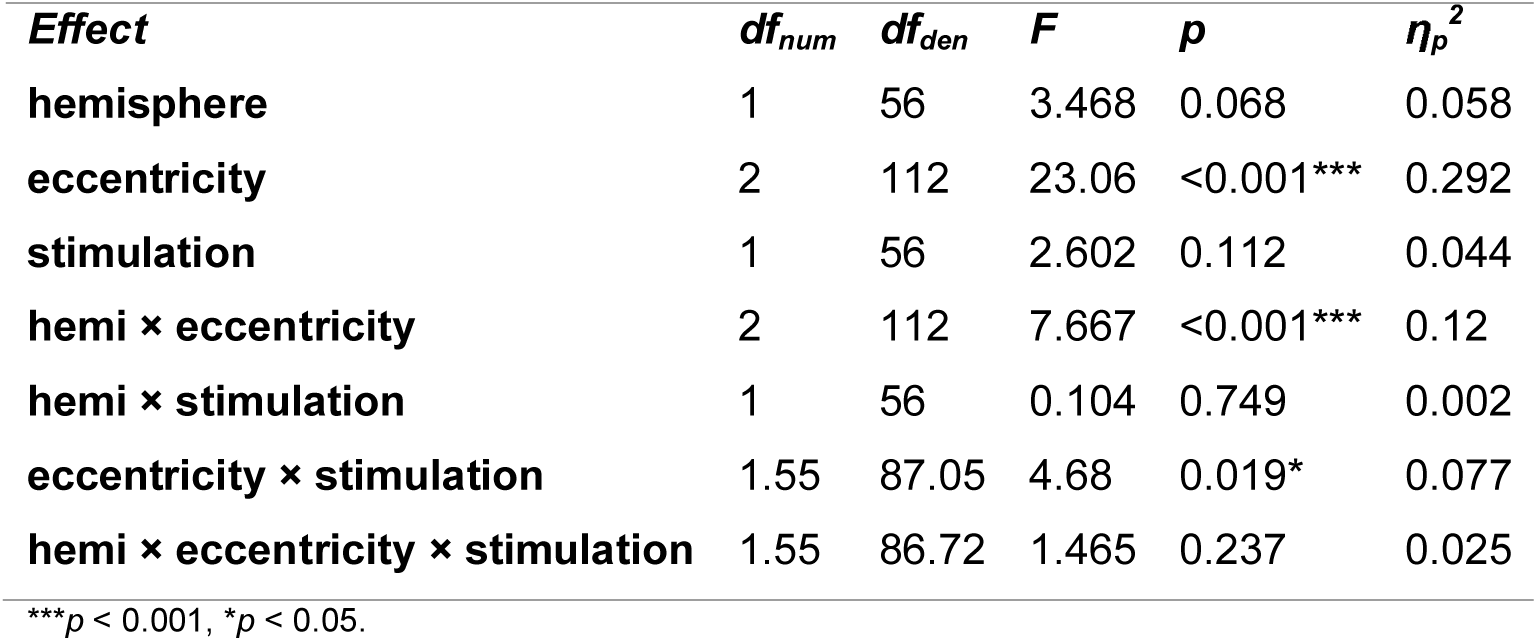
Statistical analysis of the cortical thickness effects (three-way repeated measures ANOVA)

We then tested whether this topographically specific effect was present throughout all early visual areas or exclusively in V1, which is the major cortical target of visual thalamic projections (Sincich and Horton 2005). We expressed the checkerboard effect as percent thickness change for every subject within the checkerboard representation and 1) compared these effects across areas V1-V3 with a one-way ANOVA and 2) tested the effect in each area against zero. This analysis revealed a significant effect of area (*F*(1.35, 75.39) = 3.89, *p* = 0.04, *η_p_*^2^ = 0.065), with V1 being significantly different from V2 (*t*(56) = 2.02, *p*_unc_ = 0.0489) and V3 (*t*(56) = 2.11, *p_unc_* = 0.039), albeit at uncorrected level, and V2 and V3 not differing from each other (*t*(56) = 1.08, *p_unc_* = 0.28). Furthermore, tests for a nonzero effect of checkerboard showed that a significant effect of checkerboard was present only in V1 (*t*(56) = 2.48, *p_bonf_* = 0.048, *d* = 0.33), but not in V2 (*t*(56) = 1.88, *p_unc_* = 0.065) or V3 (*t*(56) = 1.73, *p_unc_* = 0.090). The average thickness increase in V1 was 1.33 % (*SD* = 4.05), which amounted to 0.022 mm (*SD* = 0.065) in absolute units.

We further tested whether the thickness increase within the checkerboard representation of V1 was driven by the movement of the gray-white matter boundary inwards, movement of the gray-CSF boundary outwards, or both. The results of this analysis for every surface is presented in Figure 3C. There was a significant difference between the displacement of the white and the pial surface, which produced a net thickness increase (*t*(56) = 2.76, *p* = 0.008, *d* = 0.35). The displacement of the pial surface did not differ significantly from zero (*t*(56) = -0.32, *p* = 0.750), the white surface moved significantly inwards (*t*(56) = -2.11, *p* = 0.040, d = -0.279) with an average displacement of -0.029 mm (*SD* = 0.104). Similar results were obtained when the whole-brain cluster from the visual cortex was used instead of an independently defined visual ROI (white displacement: *t*(56) = -2.823, *p_bonf_*. = 0.013; pial displacement: *t*(56) = -0.200, *p* = 0.843, difference: *t*(56) = -4.168, *p* < 0.001). The thickness decrease of the frontal cluster was also driven primarily by the white surface displacement, but with an opposite sign, i.e. outward movement (white displacement: *t*(56) = 2.24, *p_unc_*. = 0.029; pial displacement: *t*(56) = 1.082, *p* = 0.284, difference: *t*(56) = 3.120, *p* = 0.002).

Finally, our additional control analysis of local changes in gray-white matter contrast showed a significant effect of hemisphere (*F*(1,56) = 20.55, *p* < 0.001, *η_p_*^2^ = 0.27), a significant effect of eccentricity (*F*(1.68, 94.23) = 35.40, *p* < 0.001, *η_p_*^2^ = 0.39), and a significant interaction between hemisphere and eccentricity (*F*(2,112) = 6.05, *p* = 0.003, *η_p_*^2^ = 0.10). Crucially, there was no interaction between eccentricity and stimulation (*F*(1.50, 84.14) = 0.11, *p* =0 .84). The full statistical results are reported in Table 3.

**Table 3.** Statistical analysis of the image intensity effects (three-way repeated measures ANOVA)

| **Effect** | **$df_{\text{num}}$** | **$df_{\text{den}}$** | **$F$** | **$p$** | **$\eta_p^2$** |
| --- | --- | --- | --- | --- | --- |
| **hemisphere** | 1 | 56 | 20.551 | < 0.001*** | 0.268 |
| **eccentricity** | 1.68 | 94.23 | 35.395 | < 0.001*** | 0.387 |
| **stimulation** | 1 | 56 | 0.158 | 0.692 | 0.003 |
| **hemi x eccentricity** | 2 | 112 | 6.049 | 0.003** | 0.097 |
| **hemi x stimulation** | 1 | 56 | 1.115 | 0.295 | 0.02 |
| **eccentricity x stimulation** | 1.5 | 84.14 | 0.113 | 0.836 | 0.002 |
| **hemi x eccentricity x stimulation** | 1.55 | 86.96 | 0.788 | 0.429 | 0.014 |

\*\*\* $p < 0.001$ , \*\* $p < 0.01$ .

### Correspondence with functional activations

To further determine how the observed changes in thickness relate to the typical BOLD fMRI effects, we compared the spatial patterns of the two effects. Comparable to the thickness effect significance peak, the fMRI significance peak was observed within the visual cortex. The fMRI significance peaks were located in the proximity of the calcarine sulcus, within the left and right lingual gyri (left MNI coordinates: x = -7.0, y = -82.7, z = -4.2; right MNI coordinates: x = 4.4, y = -82.1, z = -4.9). The two peaks in the right hemisphere were 10.13 mm apart in MNI volume space, and the two effects were overlapping (Figure 4A).

**Figure 4.**
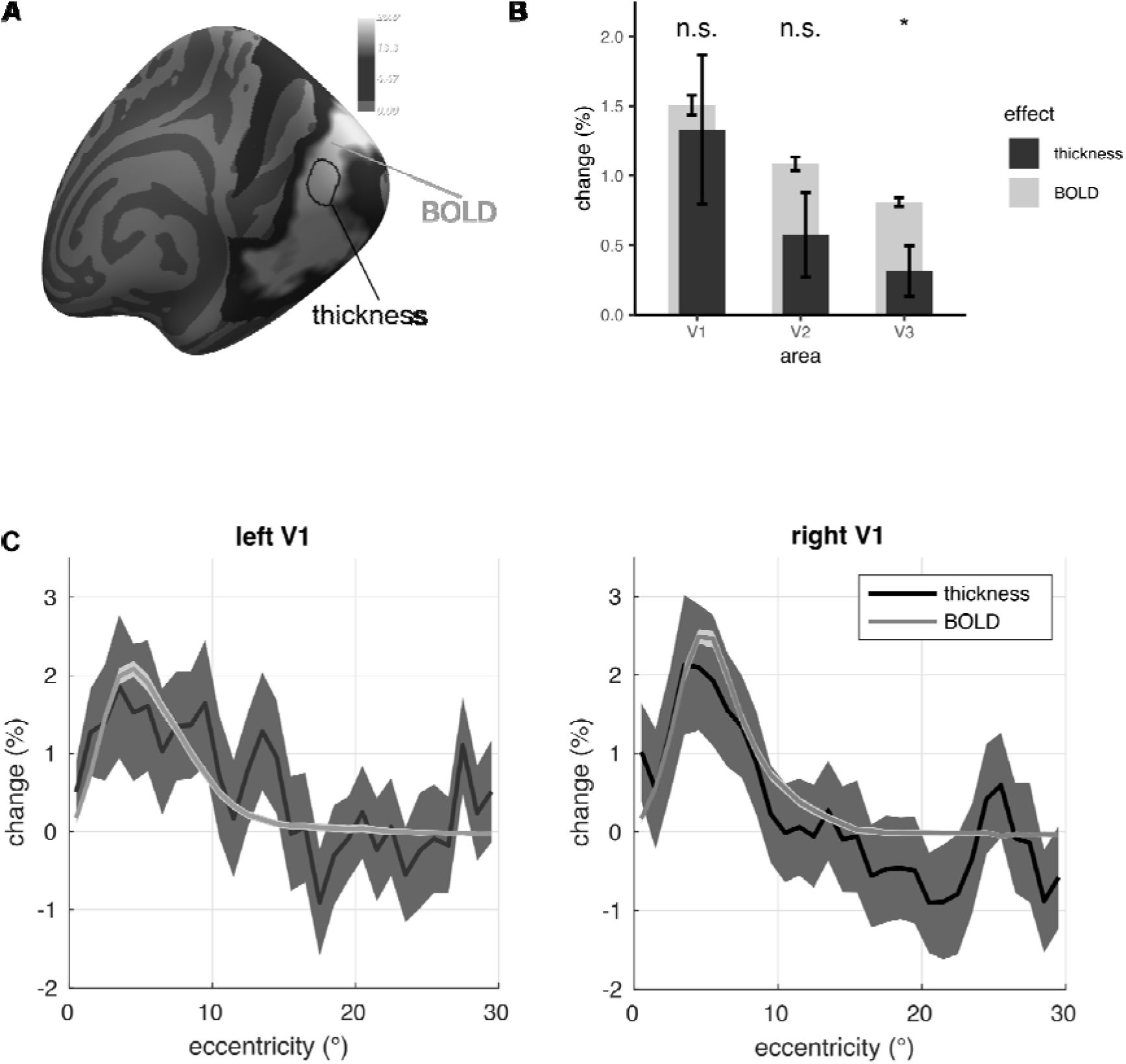
Relationship between thickness and BOLD effects. A) Overlap between whole-brain stimulation-induced thickness increase (shown as black outline) and BOLD signal (shown as overlay). Both effects are shown at p<0.01 uncorrected (for display purposes). While effects could be measured throughout the early visual cortex with fMRI, thickness changes are confined to the calcarine sulcus. B) Comparison between the magnitude of thickness change and the BOLD signal change in the ROI analysis. While there is no difference in V1 and V2, there is a significantly stronger BOLD effect in V3. C) Spatial specificity profile of both effects across 1° eccentricity bins in V1, separately for each hemisphere. Error bars and shaded error lines indicate SEM over subjects. *p<0.05 corrected.

Analysis of BOLD effects within the early visual areas revealed similarities as well as differences between the cortical thickness and the BOLD effects (Figure 4B). The two effects were similar in magnitude within the checkerboard representation in V1 (*t*(56) = 0.32, *p_unc_*. = 0.75) and in V2 (*t*(56) = 1.62, *p_unc_*. = 0.11), but diverged in V3 (*t*(56) = 2.70, *p_bonf_* = 0.03, *d* = 0.36), with the thickness effect being substantially weaker in V3. The comparison of the topographic specificity of the thickness and the BOLD changes within V1 showed a similar eccentricity profile (Figure 4C), with no differences between the two effects (all cluster-level *p* ≥ 0.64).

A more direct analysis of the relationship between thickness and BOLD effects within the checkerboard representation of V1 revealed no correlation between the two (*R* = -0.17, *p* = 0.2). There was also no significant correlation within either of the two clusters that showed significant thickness change in the whole-brain analysis (visual: *R* = -0.02, *p* = 0.90; frontal: *R* = 0.04, *p* = 0.77). Furthermore, the main effect of area in the one-way ANOVA with thickness change as a dependent variable and BOLD signal change as a covariate, remained significant (*F*(2,128.68) = 4.29, *p* = 0.016, η*_p_^2^* = 0.063).

### Simulation of the signal time course

A simple simulation of how condition order affects the detectability of an effect depending on the effect time course revealed a dissociation between the “fast” hemodynamic type of signal and a slow linear build-up (Figure 5B). There is no effect of condition order for that hemodynamic type of signal that rises within the first 5-6 seconds and subsequently reaches a plateau. However, if a signal builds up slowly within a duration of a whole scan, it is detectable in all but one condition order (1010). Interestingly, the same condition order was the only one with near-zero measured thickness estimates (Figure 5C).

## Discussion

### Summary

Our study showed that rapid increase in cortical thickness induced by checkerboard stimulation is functionally specific to the primary visual cortex, confined to the topographic representation of the stimulated visual field locations and are primarily driven by the displacement of the gray-white boundary. The topographic specificity of this effect is comparable to that of the BOLD signal within V1. In contrast to BOLD, which can be measured outside of V1, thickness changes are specific to V1. In addition, thickness changes appear to depend on the specific timing of the checkerboard presentation, suggesting a slow build-up and decay of the effect, which is incompatible with hemodynamic explanation. Altogether, our results show fast and functionally specific cortical thickness increase induced by bottom-up sensory input.

### Functional specificity of the cortical thickness effects

The functionally specific cortical thickness effects we report are largely consistent with previous studies that already demonstrated functionally specific structural changes induced by learning. For example, a study investigating cortical reorganization after balance training showed cortical thickness increases specifically within the motor cortex representations of the lower limbs and trunk, which were involved in the balance task (Taubert et al. 2016). This increase could be observed after as little as one hour of training. On a similar time scale, cortical diffusivity change has been observed within the language network for learning new lexical items (Hofstetter et al. 2017) and within the parietal cortex for an object-location association task (Brodt et al. 2018).

In a recent study, very fast (within two minutes) and functionally specific effects of motor sequence learning on cortical gray matter were reported (Olivo et al. 2022). Interestingly and in contrast to other findings, that study showed task-induced gray matter reduction instead of an increase. Furthermore, although the effects were found within the motor cortex, they did not overlap with fMRI activations evoked by the same task, suggesting that structural reorganization measured with T1-weigted imaging is distinct from task-evoked fMRI activity. We also observed some gray matter reduction in response to visual stimulation, which was localized to the superior frontal gyrus (Figure 2) and did not overlap with our fMRI activity. However, this effect was only marginally significant, and given its functionally implausible anatomical location, it may either be a false positive or an unspecific effect due to, e.g., head motion (Reuter et al. 2015).

Our primary thickness effect corresponds to an *increase* and is clearly localized within the calcarine sulcus. The core difference between our study and those discussed above is that our paradigm did not involve any form of explicit learning, which requires a complex coordination of large-scale networks under top-down voluntary control. We show that structural changes can occur even in the absence of a learning task, being driven entirely by bottom-up sensory input. Such conditions lead to gray matter enlargement and occur in regions similar to those activated in fMRI. The structural effects we observed show a high degree of functional specificity, suggesting a tight link to the locations of maximum neural activity.

Our findings are partially consistent with those of a previous study that demonstrated visually-induced structural changes in T1-weighted scans (Månsson et al. 2020). In that study, structural T1-weigthed scans were acquired while participants viewed either natural arousing images or an empty screen, observing visually induced gray matter enlargement. Surprisingly, and in contrast to our findings, areas with the most consistent structural changes in that study were located not in the primary visual cortex, as one would expect, but within the lingual and fusiform gyri. Since the comparison between viewing natural arousing images and an empty screen is rather unspecific, the resulting differences could be a product of many factors unrelated to visual stimulation, such as eye movements, attention, and arousal. In the current study, we used a luminance-matched flickering checkerboard stimulus that is known to drive the early visual areas, and an experimental paradigm that controls for fixation, attention, and arousal effects. With our paradigm we could demonstrate that thickness changes are occurring specifically within the topographic representation of the checkerboard stimulus and are strongest in V1, which is the primary target of bottom-up thalamic visual projections. We therefore demonstrated a high degree of functional specificity of the thickness increase in response to visual stimulation.

A separate analysis of white and pial surface displacement in our data revealed that the thickness changes are driven primarily by the expansion of the inner gray matter boundary further inwards. It is not clear at this stage whether this effect represents a true depth-specific change or a byproduct of volume alignment. For example, if alignment to template is driven primarily by alignment of the outer gray matter boundary and the overall brain volume and shape does not change, the local thickness differences in the visual cortex could appear as changes primarily in the inner surface. If, however, this effect reflects true depth-specific anatomical changes, it suggests an exciting possibility that gray matter enlargement may be confined to specific cortical layers (in our case to deeper layers that are closer to the white matter). Disentangling these two explanations could be a focus of future studies.

### Vascular effects or true structural reorganization

Several previous studies compared structural gray matter changes with hemodynamic effects, showing considerable overlap and suggesting a confounding effect of hemodynamics on structural T1-weighted measurements (Franklin et al. 2013; Ge et al. 2017). Despite the similarities of the thickness changes and the BOLD effects in our study, several aspects of our results support the notion that the thickness change is driven by a non-vascular mechanism. In the following, we discuss these findings in detail.

First, we could rule out that the thickness change is driven by a change in perfusion, which is typically considered a confounding factor in morphometry studies. Although cortical thickness measurements are less sensitive to changes in local image intensity compared to voxel-based morphometry studies (Chung et al. 2017), a bias related the local gray-white matter contrast (driven by variable myelin content across the cortex) on the cortical thickness estimates has been observed for this method as well, especially in the highly myelinated primary cortices (Glasser et al. 2013; Zaretskaya et al. 2018). We did not observe any change of local gray-white matter contrast which would indicate changes in intracortical T1-weighted signal intensity in our data, making changes in perfusion an unlikely explanation.

One potential alternative candidate vascular mechanism which we could not rule out is an increase in the cerebral blood volume (CBV) following neural activity (Kim and Ogawa 2012). Increase in blood volume following neural activity could lead to an overall thickness increase of a cortical sheet. If this is the case, this effect should depend on the degree of vascularization in an area. V1 is substantially more vascularized compared to V2 and V3 (Weber et al. 2008), and should therefore yield the strongest effects. Our data is consistent with the vascularization profiles of the early visual areas, with V1 showing the strongest BOLD effect, followed by V2 and V3. However, if thickness effects are driven by vasculature, we expect a covariation between the BOLD effect and the thickness effect, as both would be driven by the same underlying vascularization profile. This, however, was not the case in our data. Although similar in V1, the thickness and BOLD effects diverged in V3, with a negligible thickness effect but a comparably strong BOLD response in the latter.

Finally, our simulation of the expected effects for different condition order showed that for a hemodynamic effect, the condition order should not impact the measured thickness change. If, however, the thickness effect builds up and decays slowly, some condition sequences are more optimal for measuring the effect than others. Although our measured data for every condition order is based on few observations and should be considered with caution, it indicates that some condition sequences are less effective than others in producing a measurable thickness effect, suggesting that it builds up and decays slower than the hemodynamic response. The exact time course of the thickness effect is an important question that requires a dedicated investigation, which could be a direction for future research.

It is also worth noting that CBV changes within the gray matter can be of different origin and happen on multiple time scales. CBV increase in response to neural activity starts immediately after the stimulation onset and has a time course of a typical hemodynamic response function (Shen et al. 2008). There are, however, other plasticity-dependent CBV changes that occur on a slower time scale and parallel the true structural changes observed in animal studies. For example, CBV increase is known to accompany skill learning and has even been proposed to serve as a marker of hippocampal neurogenesis (Pereira et al. 2007). As new neurons are formed, they require better energy supply, which leads to angiogenesis. These latter CBV changes are not easily distinguishable from the tissue changes in MRI. However, CBV increase due to angiogenesis should also happen on a longer time scale than what is presented here. In this study we measured the hemodynamic response using the BOLD signal to compare the functional specificity of the thickness changes to the effects observed in a “typical” fMRI experiment. Future studies should focus on a direct comparison of CBV-weighted functional acquisitions and cortical thickness measurements using a paradigm with identical timing.

What is the fastest possible time scale of true neural tissue changes? Most animal studies investigating structural reorganization within the neocortex (e.g., dendritic spine formation) report effects on the time scale of hours and days (Holtmaat and Svoboda 2009; Caroni et al. 2012). However, non-vascular structural changes may include different processes, each of which happening at a different time scale. While dendritic sprouting, neurogenesis and angiogenesis are the longest ones and can take several days or weeks, the formation of dendritic spines can be formed on a scale of minutes to hours. Crucially, astrocyte swelling in response to neural activity can take place within seconds and could potentially account for the fast tissue changes observed in this and other studies (Macvicar et al. 2002; Johansen-Berg et al. 2012). It remains an open question whether other types of structural reorganization can happen faster than we assume, and whether these changes are sufficient to be detectable with MRI. In sum, we cannot rule out that fast cortical thickness changes due to visual stimulation are true structural changes happening on a fast time scale.

### Implications

The effects of checkerboard stimulation on cortical thickness we observed in this study appear to be rather small. Assuming a typical cortical thickness of 2 mm (Alvarez et al. 2019), they would correspond to an average increase of 1.4 % or 30 μm. Although small, these effects are on the order of those observed in clinically relevant studies. For example, in the original study of Salat et al., (2004) the cortical thickness of middle-aged adults decreased by 2.25% on average relative to the young adults. Similarly, global thickness decrease in schizophrenia is on the order of 3-4% (Guo et al. 2016). Also studies of plasticity-related thickness changes, e.g., due to meditation experience, interpreter practice or navigation training, reported effects in the same range, starting at around 2% and higher (Lazar et al. 2005; Wenger et al. 2012; Hervais-Adelman et al. 2017). In such studies, participant behavior and the stimulation environment during structural scan acquisition should be well-controlled to reduce additional variability in thickness that is due to experimental conditions.

Small effects of around 30 μm are unlikely to confound the standard fMRI studies that use gray matter thickness estimates and surface reconstruction to increase analysis precision (Dale et al. 1999; Fischl et al. 1999). They are also unlikely to confound the high-resolution "laminar” fMRI studies that typically separate cortical thickness into three depth bins, each of which corresponding to several hundred microns (Kemper et al. 2018; Polimeni et al. 2018). However, these effects should still be considered for several reasons. First, we expect some degree of inter-subject variability of the effect. Therefore, the cortical thickness increase in some subjects could be larger than 1.4 %. Second, if the cortical thickness increase linearly depends on the time of stimulation, stimulation times in a typical fMRI experiment of about 1 hour can lead to much larger effects than measured here for a 7-minute acquisition. In a scenario, in which a structural scan is acquired in the beginning of an fMRI experiment, changes in cortical thickness may lead to a gradual misalignment of the initial cortical surfaces and the subsequent functional activity. Finally, as the MRI technology develops and the human MRI is becoming capable of higher spatial resolution and spatial specificity, these effects may start playing a more significant role.

### Conclusion

In sum, we demonstrated that cortical thickness changes induced by visual stimulation are fast and functionally specific. Although the changes are extremely small and cannot confound typical fMRI, or even high-resolution “laminar” fMRI analysis, they appear to be of the order observed in typical applications of cortical thickness analysis, such as plasticity, aging or neurodegeneration studies. Future concurrent measurements of multiple aspects of hemodynamics, as well as animal and human studies investigating the time course of fast structural changes, could shed more light onto the origin of these effects.

## Acknowledgements

The authors would like to thank Thomas Zussner and Marilena Wilding for assistance with MRI data acquisition, Magdalena Lhotka for help with participant recruitment and Elias Kerschenbauer for proof-reading. We also thank Jonathan Polimeni for helpful discussion. This work was funded by the BioTechMed-Graz, Austria (Young Research Group Grant to N.Z.) and by the Institute of Psychology of the University of Graz.

## Author contributions

Natalia Zaretskaya: Conceptualization, Methodology, Software, Formal analysis, Investigation, Writing - Original Draft, Visualization, Funding acquisition

Erik Fink: Investigation, Formal analysis, Writing - Review & Editing, Project administration

Ana Arsenovic: Software, Validation, Writing - Review & Editing

Anja Ischebeck: Validation, Resources, Writing - Review & Editing

## Footnotes

1 https://cfn.upenn.edu/aguirre/wiki/doku.php?id=public:retinotopy_template

2 https://www.mathworks.com/matlabcentral/fileexchange/71737-permutest

