## Supplemental Figure S1 for "Fast and functionally specific cortical thickness changes induced by visual stimulation"

### Title:

Universitaetsplatz 2

8010 Graz

Austria

**Supplementary Figure S1**

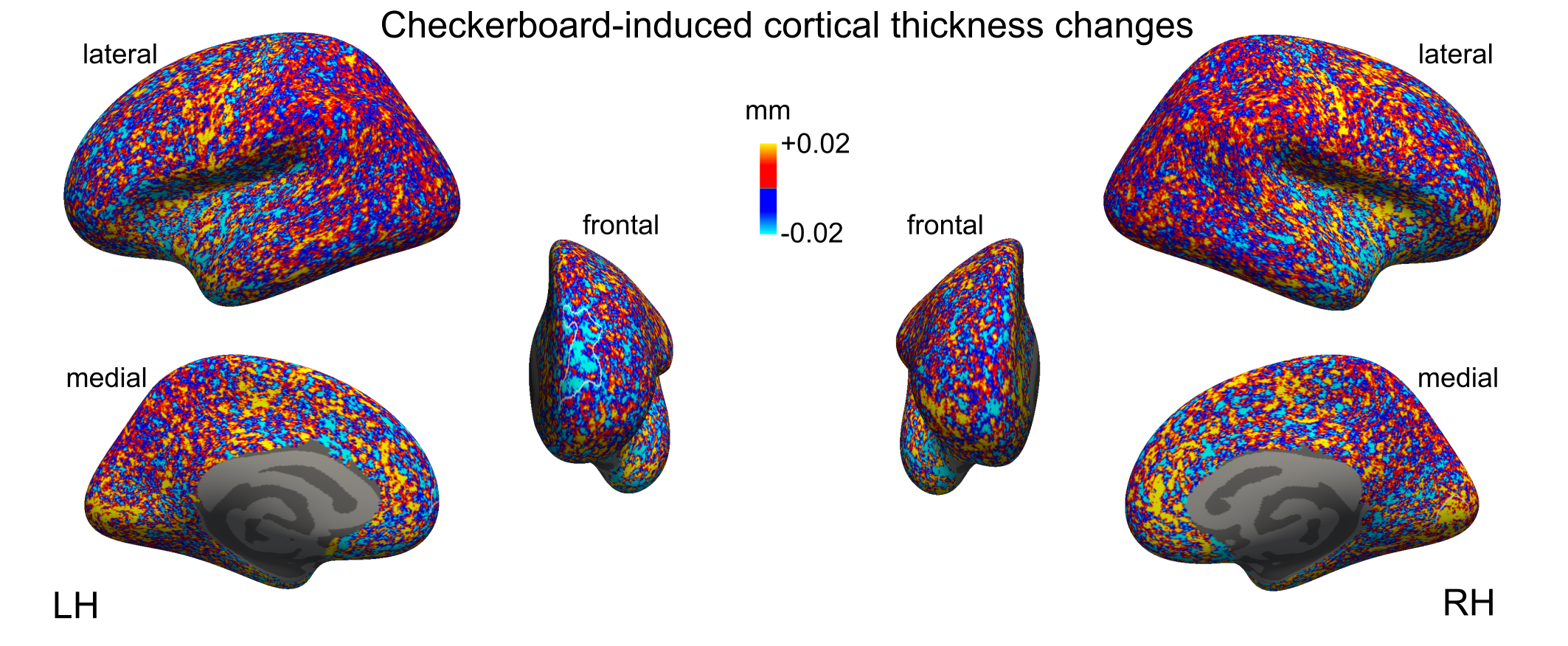

Figure S1 (related to Figure 2). Whole-brain cortical thickness changes induced by checkerboard stimulation. Maps represent average difference between “stimulation on” and “stimulation off” conditions in mm at every surface vertex in the same way as Figure 2, but prior to smoothing.
